# Oomycetes during 120,000 years of temperate rainforest ecosystem development

**DOI:** 10.1101/042341

**Authors:** Ian A. Dickie, Angela M. Wakelin, Laura Martinez-Garcia, Sarah J. Richardson, Andreas Makiola, Jason M. Tylianakis

## Abstract

The occurrence of plant-associated oomycetes in natural ecosystems and particularly during long-term ecosystem development is largely unknown, despite the importance of many oomycetes as plant pathogens. Using DNA sequencing from roots, we investigated the frequency and host relationships of plant-associated oomycete communities along a 120 000 year glacial chronosequence, comprising site ages with rapid compositional change (“early succession”; 5, 15, and 70 years old soil); relatively stable higher-diversity sites (“mature”, 280, 500, 1000, 5000, 12000 years); and ancient, nutrient-limited soils with declining plant diversity and stature (“retrogression”, 60 000, 120 000 years). Plant-associated oomycetes were frequent in all three early successional sites, occurring in 38 – 65% of plant roots, but rare (average 3%) in all older ecosystems. Oomycetes were highly host specific, and more frequent on those plant species that declined most strongly in abundance between ecosystem ages. The results support the particular importance of plant-associated oomycetes in early succession (up to 70 years). High host specificity and correlations of abundance of oomycete inside roots with declining plant species are consistent with oomycete-driven successional change.

## Introduction

As ecosystems develop, biotic communities undergo succession both in composition and in the nature of mutualistic and antagonistic interactions. Many of these changes have been well studied, including changes in plant communities and traits, competition intensity, and mutualisms (Lambers et al, 2008). Nonetheless, long-term successional changes in composition and community interactions of many other organisms, such as oomycetes, remain poorly understood. Indeed, oomycetes in natural ecosystems are often out-of-sight out-of-mind, until large-scale forest diebacks occur (Scott & Williams, 2014). This lack of information is particularly apparent in forest disease outbreaks, where it is often unclear whether the oomycete is native, from where it may have originated, or what background levels of oomycetes are typical in healthy ecosystems (Podger & Newhook, 1971). Oomycetes occur as part of healthy ecosystems and play important roles in ecosystem processes (Castello et al, 1995; Gómez-Aparicio et al, 2012; Bever et al, 2015; Geisen et al, 2015). For example, *Pythium* and other oomycetes (Oomycota = Peronosporomycota) can drive negative density dependence in plants (Packer & Clay, 2000) and hence are believed to contribute to the maintenance of forest plant diversity (Mangan et al, 2010a). Oomycetes are also an important component of soil biodiversity, as pathogens of plant roots and soil invertebrates, and as saprotrophs (Arcate et al, 2006; Geisen et al, 2015). One way to better understand the role of oomycetes in natural ecosystems is to look at gradients of ecosystem development, which have a long history of providing insight into plant communities.

A useful way to consider ecosystem development, at least from a plant-community perspective, is based on compositional change through three phases: early succession, mature, and retrogression. Early succession (both primary and secondary) can be defined as the period in which communities undergo major compositional change. In the context of plant communities, this means that early successional plants are unlikely to regenerate under conspecifics and are instead adapted for dispersal into new sites. Mature ecosystems can be defined by compositional stability over timespans greater than one generation, such that species are regenerating in the presence of established conspecifics (Veblen, 1992). Finally, over very long time periods ecosystem retrogression eventually develops, where the progressive loss of nutrients leads to a decline in plant diversity and a loss of biomass and canopy stature (Peltzer et al. 2010). This shift to retrogression entails slow compositional change, favouring species with life history traits that are adapted to stressful, low-nutrient environments.

Although oomycetes can be saprotrophic, they have been proposed to play particularly important roles at all stages of ecosystem development as potential plant pathogens. One view of soil pathogens in relation to ecosystem development is that they play a critical role in driving compositional change in early succession (Van der Putten et al, 1993; Van Der Putten, 2003). This view is based, in part, on the assumption that newly establishing plant species will encounter only low levels of pathogens in the absence of conspecific host plants during succession. Over time, these established plants accumulate pathogens, which prevent establishment of conspecific seedlings but have less negative effects on seedlings of other plant species. Part of the role of pathogens in succession is driven by plant age, as pathogens that kill seedlings are often tolerated by established plants (Martin & Loper, 1999; Van Der Putten, 2003). This implies that early-successional (*r*-selected) plants may have limited selection for pathogen resistance. Instead, these species may accumulate and tolerate pathogens as mature plants, and rely on seed dispersal into new habitats to escape pathogens during the vulnerable seedling stage (Van Der Putten, 2003). Support for the role of antagonists in early succession comes from studies of invertebrate antagonists in grasslands, where dominant early-successional plants are more suppressed than subordinate or late-successional species (De Deyn et al, 2003) and from studies showing negative plant-soil-feedback early in succession (Kardol et al, 2006). Oomycete pathogens have also been shown to be important in the mortality of the early-successional tree *Prunus serotina* (Packer & Clay, 2000; Packer & Clay, 2003). These examples span from short-term grassland to decadal scale forest patterns, suggesting a widely applicable ecological pattern.

While much evidence supports the role of pathogens in early succession, there are also suggestions that pathogens, in general, increase in mature ecosystems (Reynolds et al, 2003; Bardgett et al, 2005; Peltzer et al, 2010). Soil biotic communities as a whole increase in biomass and species diversity over periods of tens to hundreds of years (Bardgett et al, 2005), and pathogen communities may follow similar patterns of increasing biomass and diversity. Further, plant competition is high in mature ecosystems, and Gilbert (2002) suggests that competition may stress plants and increase susceptibility to pathogens. Pathogen host-specificity in these systems leads to density-dependent mortality (Augspurger, 1984). Evidence for negative density dependence has come from both tropical and temperate forests (Packer & Clay, 2003; Mangan et al, 2010b), although other studies have found positive density dependence or no effect (Reinhart et al, 2012).

Finally, a recently proposed third view of the role of pathogens in ecosystem development is that soil pathogens may be particularly critical in retrogressive ecosystems (Laliberté et al, 2015), where declining soil nutrient availability results in declining vegetation stature and diversity (Richardson et al, 2004; Porder et al, 2007; Peltzer et al, 2010). Laliberté et al. (2015) suggest that the highly weathered, P-limited soils of retrogressive ecosystems favour species with ephemeral roots to maximize nutrient uptake, but that this imposes a trade-off with increased susceptibility to pathogens. Ecosystem retrogression has been linked to increased specific root length (length per unit mass), thinner roots, and increased root branching (Holdaway et al, 2011), all of which may increase susceptibility to root pathogens (Laliberté et al, 2015).

Based on the three views from the literature, we tested the non-exclusive hypotheses that oomycetes would be associated with (1) early-successional, (2) mature, and/or (3) retrogressive ecosystems. We used an extensively studied soil and ecosystem development chronosequence created by the Franz Josef glacier in New Zealand, where glacial advances and retreats have created a series of soils of varying age in close proximity (Walker & Syers, 1976; Richardson et al, 2004), allowing space-for-time substitution as an approach to infer change over long time periods (Pickett, 1989). We focussed on oomycetes in living plant roots, in order to link oomycete communities to particular hosts. The presence of an oomycete in roots of specific plant species does not necessarily indicate that it causes disease in that plant. Indeed, mature plants may often tolerate organisms that would kill conspecific seedlings. It is also conceivable that oomycetes are living within the roots of plants on which they would never cause disease, regardless of plant age. Nonetheless, while the presence of oomycetes does not necessarily indicate disease, the absence of oomycetes from a plant can be taken as an indication of absence of oomycete-driven disease.

## Materials and Methods

### Study site and sampling

The Franz Josef chronosequence includes early-successional (5, 15, 70 years of development), mature (280, 500, 1000, 5000, 12 000 years), and retrogressive (60 000, 120 000 years) sites. As soils age, there are dramatic changes in nutrient availability (declining P, increasing and then decreasing N), pH, and physical properties (Walker & Syers, 1976). These changes are linked to changes in plant communities, with plant biomass increasing through succession to mature stages, and then entering retrogression where declining soil nutrients result in a concomitant decline in plant biomass, stature, and diversity (Richardson et al, 2004) and shifts in root traits (Holdaway et al, 2011). Vegetation shifts from sparse herbaceous plants and sub-shrubs (5 years) to shrub land (15 years) to angiosperm forest (70 years), followed by an increasing dominance of large gymnosperm trees (Podocarpaceae) through mature stages, with an eventual decline in plant biomass, canopy height, and canopy closure in retrogression. Rainfall is high along the entire chronosequence, ranging from 3.5 to 6.5 m, and all sites are below 365 m elevation (Richardson et al, 2004).

In a previous study, we collected 510 individual plant roots from ten sites along the chronosequence and characterised both plant identity and arbuscular mycorrhizal fungal communities in those roots (Martínez-García et al, 2015). Here we use these same samples but used taxon-specific primers to amplify DNA from oomycetes. The collection of samples is described in full in Martinez-García *et al.* (2015). In brief, we collected 51 or 52 (at two sites) individual fine root (< 2 mm diameter) fragments, taking a single root fragment (approx. 15 mg dry weight) every 2 m along two parallel transects with equal sampling from three depths (up to 20 cm depth max). This depth is likely representative of the established roots a seedling might encounter during the first year of growth and could be sampled in a standardised way at even our youngest sites, but may omit deeper roots in older sites. For the 5-year-old site, which comprised sparse vegetation in rocks, most soil was root free. We therefore collected the nearest plant to the sample point and sampled one root from that plant. Roots from all sites were thoroughly rinsed in water and dried before DNA extraction. DNA was extracted using a MoBio soil DNA kit, and plant species identified by PCR amplification and DNA sequencing of the tRNL gene region, except for the 5-year-old site, where the plant was already known due to collection method. The one difference in the sampling between this study and the prior study (Martínez-García et al, 2015) was that in the earlier paper a spare (” B”) sample was used if the first (” A”) sample failed to yield both plant and arbuscular-mycorrhizal fungal PCR products. In the present study we did not use the “B” samples, as it would have made quantification difficult.

### Oomycete PCR and identification

We primarily based our oomycete detection, operational taxonomic unit (OTU) clustering, and identification on a nested PCR of large-subunit (28S) DNA (supplementary methods). Initial results using T-RFLP suggested that most samples contained either no oomycetes or only a single OTU, hence direct sequencing was used for identification. A negative (omitting DNA template) and a positive control (including genomic DNA from a *Pythium* species) were included in every PCR. PCR products were purified using DNA Clean and Concentrator Kit (Zymo Research Corporation) prior to performing Sanger sequencing (Canterbury Sequencing and Genotyping, University of Canterbury, New Zealand).

Some ecosystem ages had a high proportion of samples that failed to amplify a product of the expected size in the first-round PCR. In order to ascertain whether these were true absences of oomycetes or due to PCR inhibition, nine samples from each of these sites with the highest failure rate (500, 100, 1200, 60 000 and 120 000 yrs) were tested for PCR inhibitors by repeating PCR reactions in duplicate for each sample, with one of the duplicate reactions spiked with 20 ng positive control DNA. Only one out of the 45 samples spiked with positive control DNA failed to amplify, suggesting PCR inhibitors were not likely to be causing low detection rates.

To confirm identities of sequenced oomycetes, samples that produced positive large subunit PCR products were sequenced for the internal transcribed spacer (ITS) 1 region (supplementary material). As DNA sequences were obtained from environmental samples (root fragments) potentially containing multiple oomycetes, the ITS sequence may or may not represent the same species as the 28S sequence. ITS sequences were therefore used to help inform identification, but frequency analyses were based on the 28S results.

#### Vegetation cover

The percent cover of all vascular plants within sites was measured following the standard relevé (or ‘ recce’) method for describing New Zealand vegetation (Hurst & Allen, 2007) in a 20 x 20 m area located within 50 m of the root sample transects. For testing the correlation of plant cover change with oomycete frequency we used the mid-point of vegetation cover classes summed across height tiers as a measure of the abundance of each plant species.

#### Sequence bioinformatics

All DNA sequences were matched against GenBank using BLAST to find the closest matching sequence, and, where the closest match had no reliable taxonomic identity, the closest matching sequence associated with a taxonomic identity. Sequences that matched non-oomycete specimens were discarded. Three sequences had a closest match to a *Spongospora* (Cercozoa) sequence, but only at 78% identity. We therefore considered these three sequences to likely be root-associated protists (Bulman & Braselton, 2014) and retained them in the analysis. Out of 510 root samples, 122 had positive PCR products with oomycete primers. After filtering for sequence quality and matching to an oomycete sequence, the final dataset contained 91 DNA sequences of oomycetes with a mean sequence length of 310 bp. Plant IDs were obtained for 458 samples overall and 86 of the 91 samples with oomycete products through DNA sequencing of the trnL gene (Martínez-García et al, 2015). We clustered sequences into OTUs using BLASTn to merge any sequences with > 97% similarity over at least 95% of the shorter sequence length based on single-linkage clustering. This allowed us to form species concepts for testing host specificity, resolved some matches to “uncultured oomycete” and suggested where the gene region used lacked sufficient resolution to distinguish species.

#### Statistics

Most of our analyses are based on the frequency of roots with oomycetes, testing the prevalence of oomycetes within different plant species and sites, rather than quantifying abundance within any particular root fragment. Changes in the frequency of oomycetes as a function of successional stage were tested by treating age as a three-level factor. Sites were allocated to successional stage levels on the basis of soil nutrient concentrations, vegetation height, species richness and biomass, and plant traits (Walker & Syers, 1976; Richardson et al, 2004). Early-successional sites (5, 15, 70 years) had abundant N-fixing trees; vascular plant traits of high foliar nutrient concentrations, low leaf mass per unit area; very high soil P, rapidly accumulating biomass and species richness of vascular plants, and increasing nitrate-N. Mature ecosystems (280 to 12 000 years) had slow rates of change in species richness, height and biomass among sites. These sites also had the highest biomass and plant diversity. Retrogressive sites (60 000, 120 000 years) had a decline in vegetation height, richness and biomass and exceptionally proficient phosphorus resorption (Richardson et al, 2005; Peltzer et al, 2010; Vitousek et al, 2010).

We tested changes in the frequency of oomycetes as a function of successional stage as a factor and log-transformed ecosystem age as a linear variate using a binomial glm, fitted with quasi-likelihood to account for over-dispersion. The difference between the successional stage and ecosystem age models was tested using analysis of deviance, using an F test as recommended for quasibinomial fit. For these analyses n = 10 based on 512 individual samples.

We wanted to test whether declining plant species had higher than random oomycete levels, as would be expected if oomycetes are an important driver of community changes with ecosystem development. To do so, we calculated the change in cover of each plant species from one time point (t_0_) to the next (t_1_) as a log ratio (log((cover t_0_ +1) / (cover t_1_+1))) and then used a binomial mixed effects model to test whether the presence / absence of oomycetes in root fragments at t_0_ could be predicted by change in cover from t_0_ to t_1_ across the nine intervals between site ages. This test was carried out using lmer in the R package lme4, with site age and plant species (to account for species occurring across multiple sites) as random effects and the canonical logit link function. Host-specificity was tested within ecosystem age using chi-square tests.

## Results

### Oomycete frequency as a function of stage of ecosystem development and ecosystem age

In early-successional ecosystems (5, 15 and 70 years), 38 to 65% of root samples had oomycete DNA detected (Figure 1). In contrast, in mature and retrogressive ecosystems (280 to 120 000 years) an average of only 2% and never more than 8% of roots had oomycete DNA found (Figure 1). Treating the sites as representing three stages, the early successional stages had a significantly higher frequency of oomycetes than mature stages (quasibinomial family glm; *t* = −6.6, *P* = 0.00050) or retrogressive stages (*t* = −4.6, *P* = 0.0025), and the difference between mature and retrogressive ecosystems was not significant (*t* = −1.1, *P* = 0.32).

**Figure 1.**
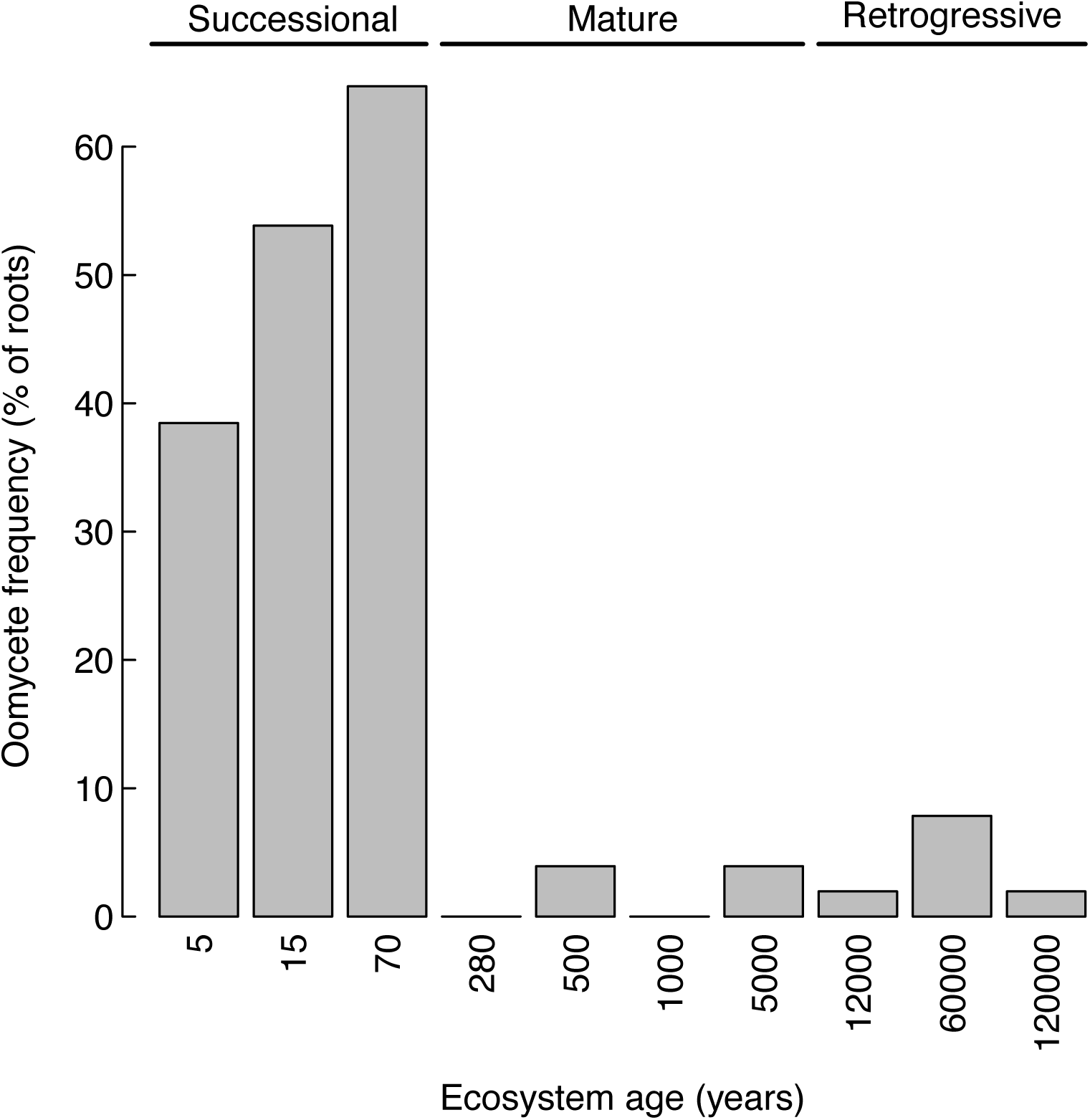
Frequency of oomycetes in plant roots (%) as a function of ecosystem age. The 5, 15, and 70-year-old sites represent a successional sere from rock field through shrubland to angiosperm forest. Sites 280 to 5000 years of age are mature forest with high biomass and plant diversity, while after 12 000 to 60 000 years plant biomass, stature, and diversity decline due to retrogressive P-limitation.

Oomycete frequency was also significantly related to log-transformed ecosystem age (quasibinomial family glm; *t* = −2.75, *P* = 0.025). While statistically significant, this relationship was a poor fit to the observed pattern of an increase for the first 3 site ages followed by uniformly low levels of oomycetes at all subsequent site ages (Figure 1). Analysis of deviance strongly supported the ecosystem-stage categorical model over the linear ecosystem-age model (*F* = 35.11, *P* = 0.00058).

Those plant species that showed the largest decline in cover between ecosystem ages also had the highest frequency of oomycetes in their roots (Figure 2, *z* = −2.4, *P* = 0.018).

**Figure 2.**
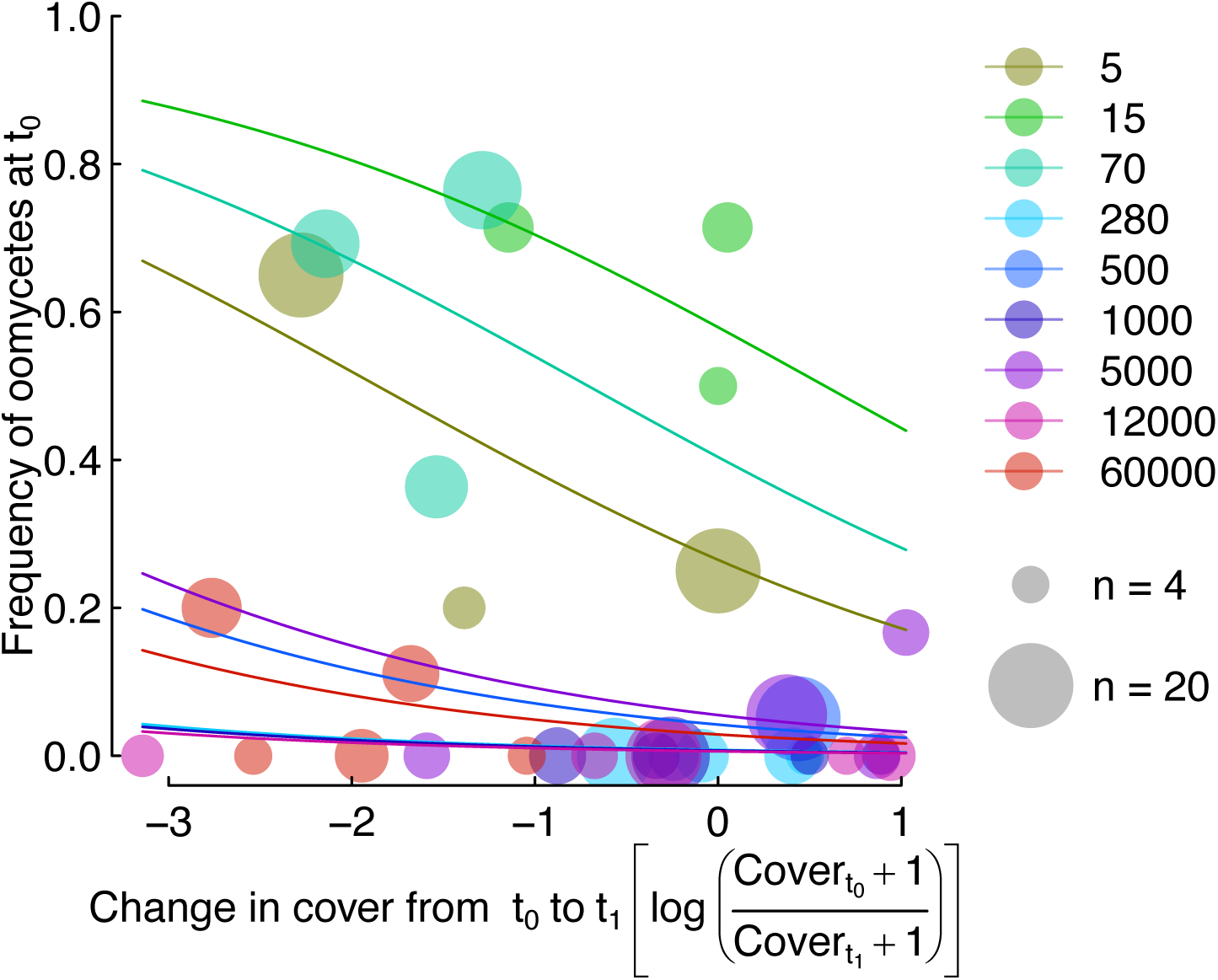
Oomycete frequency as a function of vegetation decline with circles representing each plant species between site transitions. Points scaled by number of samples for each plant species within site. Lines indicate random effects to account for site in a binary mixed effects model with random intercepts for site and plant species. Colour coding in reflects the site age at t_0_ for each transition, hence no points for 120 000 years are shown.

### Oomycete diversity, host specificity and ecosystem age

A total of 37 different OTUs were found, with 10 OTUs found more than once (Table 1). The most frequent OTU occurred 17 times, and had DNA-sequence affinities to *Lagena radicicola* and an uncultured oomycete from New York, USA agricultural soils (Table 1).

**Table 1.**
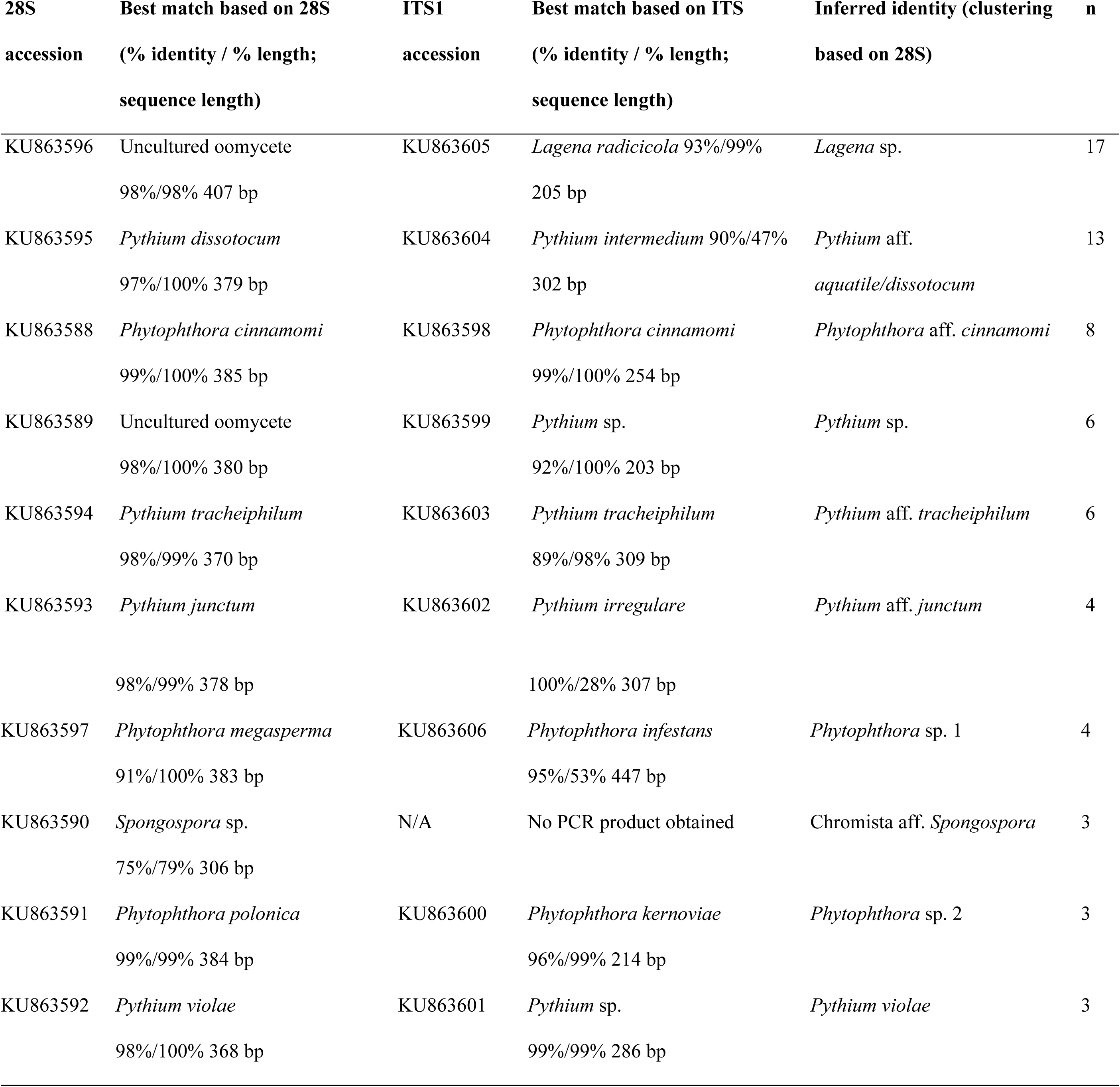
Identity of non-singleton OTUs as clustered based on 28S, with accession numbers and best matching 28S and best matching ITS1 sequences, the inferred identity, and the number of sequences in the OTU.

Within ecosystem ages, there was a significantly non-random distribution of oomycete OTU identity across plant species at 5 (5 oomycete OTUs with 20 occurrences across 4 plant species, χ^2^ = 42.96, df = 12, p-value = 0.000022) and 15 years (15 oomycete OTUs, 28 occurrences, 14 plant species, χ^2^ = 225.87, df = 182, p-value = 0.015), but only marginally significant at 70 years (16 oomycete OTUs, 30 occurrences, 7 plant species, χ^2^ = 109.85, df = 90, p-value = 0.076; Figure 3). There were too few observations at the ecosystem stages after 70 years to meaningfully test for plant by OTU host-specificity at later ecosystem stages.

**Figure 3.**
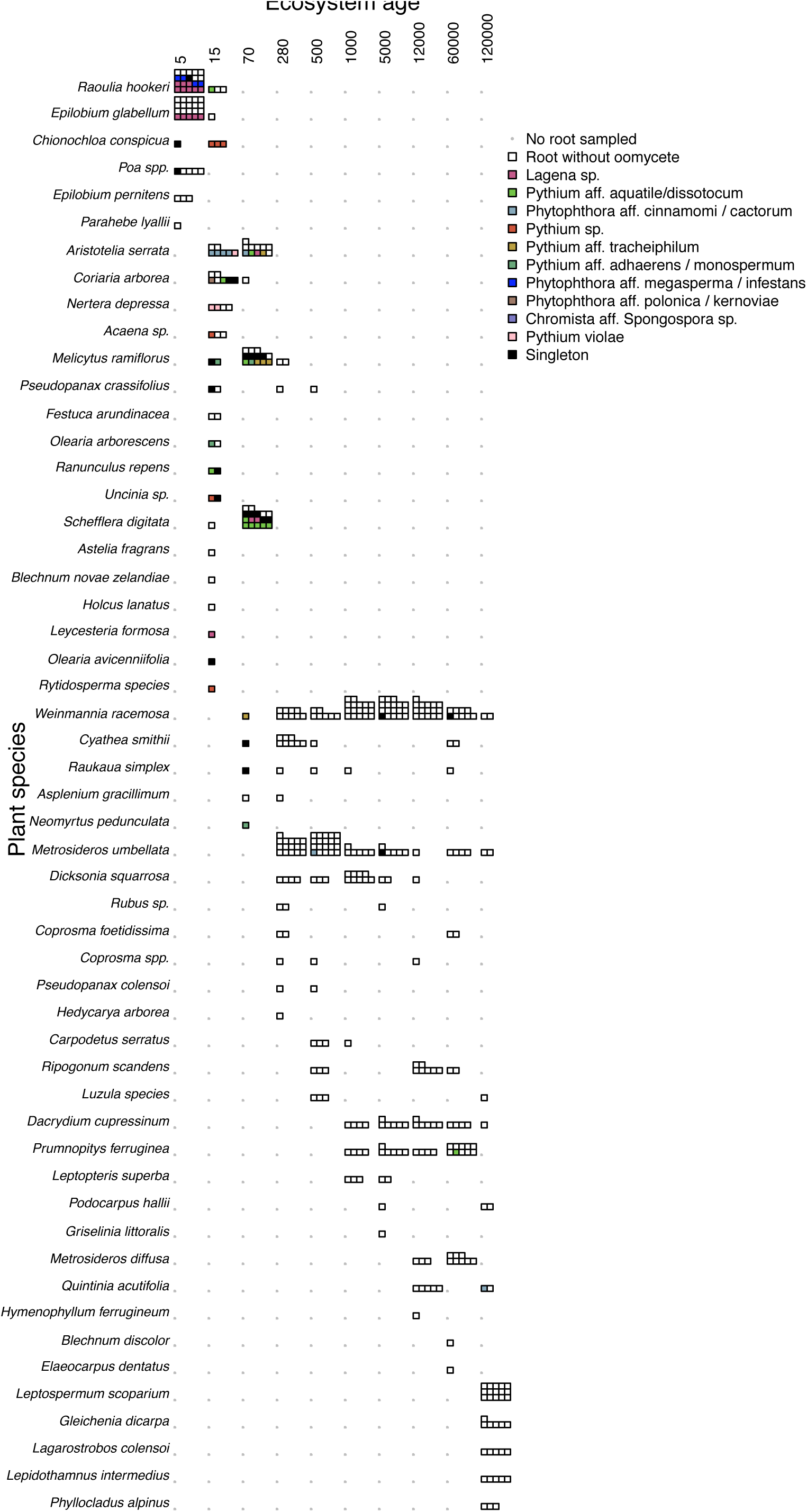
Plant by site age visualisation of oomycete frequency within sampled roots. Each square represents a single root sample, with coloured squares indicating the presence of oomycete pathogens with colours indicating OTU and black indicating an OTU found only once. See Table 1 for OTU information. Grey dots indicate species not found in root samples for that site.

## Discussion

Our results strongly support the importance of ecosystem age as a determinant of oomycete communities in plant roots, with high frequencies of oomycetes observed in the three earliest ecosystem stages (5, 15, and 70 years) but only very low levels thereafter. Although the importance of oomycetes as pathogens is often discussed in the ecological literature (Bagchi et al, 2014), relatively few studies have actually quantified oomycete frequency in natural ecosystems (Gómez-Aparicio et al, 2012). Further, we believe our results are the first study of oomycetes throughout a complete ecosystem development sequence. The community of oomycetes was diverse, including multiple *Pythium* and *Phytophthora* species and, surprisingly, a very frequent sequence matching *Lagena* on two small sub-shrubs: *Raolia hookeri* (Asteraceae) and *Epilobium glabellum* (Onograceae). *Lagena* has previously only been reported as a pathogen on grass (Blackwell, 2011).

In terrestrial ecosystems, oomycetes occur as pathogens and commensals of plants and other eukaryotes and, less commonly, as saprotrophs (Marano et al, 2016). By sampling plant roots, our results focussed primarily on endophytes. Our observation of high host-specificity is also consistent with a biotrophic interaction and the correlation of oomycete frequency with subsequent plant decline is consistent with pathogenicity. However, while we believe that the oomycetes we observed were primarily biotrophic, the gene regions we sequenced lacked sufficient resolution to rule out the possibility that some of the *Phytophthora* we observed could be in clade 9, which contains some known saprotrophs (Marano et al, 2016). The other clades of oomycetes where saprotrophy is known to occur (Marano et al, 2016) were not observed.

### Ecological role of oomycetes in plant roots in early successional ecosystems

It has been suggested that pathogens, including oomycetes, drive early-successional change in plant communities (Kardol et al, 2006). The high frequency of oomycetes in early successional ecosystems is consistent with this hypothesis. In addition, those plant species that had the highest frequency of oomycetes at a given ecosystem age declined in percent cover most strongly before the next age. The higher frequency of oomycetes on plant roots in early-successional ecosystems compared to mature and retrogressive ecosystems was not due to PCR inhibition in older sites, as samples spiked with positive control oomycete DNA showed no evidence of inhibition.

Oomycetes, particularly *Pythium*, are often tolerated as endophytes by established plants, but prevent seedling establishment of the same plant species (Martin & Loper, 1999; Van Der Putten, 2003). On that basis, we do not argue that the observed oomycetes were necessarily having any direct negative effect on established plants. Instead, we suggest that the correlation of oomycete frequency with decline in cover between ecosystem ages is congruent with the suggestion that these pathogens are primarily preventing re-establishment of plants and hence contributing to vegetation change across cohorts (Van Der Putten, 2003).

Our definition of “early succession” comprises the major transition from rock field (5 years) to shrub land (15 years) to forest (70 years). Therefore, our finding of high pathogen levels is potentially consistent with Packer & Clay (2000), who found strong negative feedback in *Prunus serotina* in a forest described as “at least 70 years old” (Packer & Clay, 2000), despite the differences between our primary succession and what was likely a secondary succession in the Packer & Clay study.

We believe most of the change in oomycete frequency across ecosystem stages reflects changes in plant species composition. While oomycete frequency was significantly correlated with site age as a linear predictor, treating sites as three distinct stages resulted in a much stronger model fit. Nonetheless, changes in oomycete frequency may also, in part, reflect a substantial site-age effect and direct effects of changing soil environments on oomycete populations (akin to the “habitat” hypothesis for mycorrhizal fungi of Zobel & Opik 2014). Meaningful testing of soil variables as predictors of oomycetes was not possible, as soil properties were strongly correlated with soil age.

### Absence of oomycetes in plant roots in older ecosystems

It has been suggested that oomycetes are important in mature (Bagchi et al, 2010; Bagchi et al, 2014) and retrogressive ecosystems. While Packer and Clay (2000) clearly demonstrated oomycetes determined seedling dynamics in a relatively young temperate forest, a more recent study by Bagchi and colleagues (2014) in a tropical seasonal forest found that eliminating fungi alone had a stronger effect on negative density-dependence than eliminating both fungi and oomycetes. Laliberté et al. (2015) suggested that root traits adapted to highly P-limited retrogressive ecosystems would increase susceptibility to root pathogens, and that oomycetes could drive plant diversity in these ecosystems (Albornoz et al, 2017).

Our results do not support an important role of root associated oomycetes in this retrogressive sequence, which contrasts with the recent findings from a western Australian system where oomycetes play key roles in older ecosystems (Albornoz et al, 2017). In part this may reflect the importance of non-mycorrhizal Proteaceae in Australia. The very fine roots of the Proteaceae may make this plant family particularly susceptible to pathogens (Laliberté et al, 2015). Further comparative studies across multiple retrogressive ecosystems are needed to resolve the circumstance under which each pattern is more common.

### Novel organisms or native?

Having baseline data on oomycete ecology may be important for understanding and managing future pathogen outbreaks. For example, *Phytophthora agathidicida* (= *Phytophthora* taxon *Agathis*) has been implicated as a cause of forest dieback of the iconic New Zealand-endemic *Agathis australis*, but, like many other oomycete disease outbreaks, the origin of *Phytophthora agathidicida* remains uncertain (Weir et al, 2015). The single-direction sequencing we performed directly from environmental samples was not designed for accurate phylogenetic classification, even with two gene regions sequenced per sample, but it does suggest considerable diversity of *Phytophthora*. We found sequences with affinities to a number of species already known to be present in New Zealand, including *P. cinnamomi*, *P. cactorum*, *P. infestans* and *P. kernoviae* (Scott & Williams, 2014). Given that our samples came from relatively pristine ecosystems, we believe these OTUs are likely to be from native *Phytophthora*.

More surprising was that the most common OTU in our data had affinities to *Lagena radicicola* in both 28S and many of the ITS sequence results. *Lagena* is presently considered to be a monotypic genus widespread as a pathogen of grasses in North America (Blackwell, 2011), but Barr and Désaulniers (1990) suggest it may be under-reported due to being morphologically very similar to *Pythium*, and not being easily culturable. Spores described as “resembling *Lagenocystis* [syn. *Lagena*] spp.” were noted by Skipp and Christensen (1989) in New Zealand *Lolium perenne* pastures, but our finding is the first report of a putative *Lagena* species outside of grasses. We believe this most likely reflects a lack of prior knowledge, as very few prior studies have used molecular methods to detect oomycetes from healthy forest ecosystems, and none of those have taken place in temperate southern hemisphere rainforest.

## Conclusions

Much of the Earth’ s surface is covered in young, early successional ecosystems comparable to our early successional sites of 5, 15, and 70 years (Vitousek et al, 1997; Haddad et al, 2015). The strong plant-host specificity and correlation of plant decline and oomycete frequency is consistent with the suggestion that these oomycetes as pathogens contribute to vegetation succession. While we do not find frequent oomycetes in older ecosystems, oomycete DNA was detected in all but one site. These low levels of oomycetes in roots, along with oospores in soil, may still be sufficient to prevent establishment of susceptible plant species or to drive pathogen outbreaks following external climatic stressors, but seem incompatible with density dependent mortality being driven by oomycetes on established plant roots. More broadly, our results support the concept that oomycetes are a diverse part of native ecosystems. Understanding this native diversity in is critical to better understand, prevent, and manage future oomycete disease outbreaks (Dickie et al, 2017).

## Acknowledgements.

We thank D. Peltzer for help with locating plots, characterising vegetation, and editorial input, R. Holdaway for helping with data and other advice, H. Ridgway for providing positive control DNA, N. Merrick for sequencing, R. Buxton and C. Morse for collecting additional plant trait data, and S. Wakelin, L. Condron, J. Bufford, S. Bulman and the Mycorum discussion group at Lincoln University who provided valued advice and input. Funding was provided by Tertiary Education Commission funding for the Bio-Protection Research Centre (I.A.D., AMW, JMT), Lincoln University Research Fund (IAD, AMW), a Rutherford Fellowship (JMT), and Core Funding to Crown Research Institutes (LMG, SR).

## Data Accessibility

- DNA reference sequences: GenBank (accession numbers in Table 1)

- Vegetation data are archived in the NewZealand National Vegetation Survey Databank (<u>https://nvs.landcareresearch.co.nz</u>; last accessed 8 Feb 2016).

Supplemental files S1: Detailed molecular methods and discussion.

